# Somatic Hdac4-902fs mutations lead to loss of HDAC4 function through nonsense-mediated mRNA degradation

**DOI:** 10.1101/2025.10.14.682279

**Authors:** Harikrishnareddy Paluvai, Mark E. Pepin, Friederike Schreiter, Alireza Saadatmand, Marco Hagenmueller, Johannes Backs

## Abstract

Histone deacetylases (HDACs) are essential chromatin regulators and are involved in the regulation of gene expression by removing acetyl groups from histone and non-histone proteins. Histone deacetylase 4 (HDAC4) is known to regulate the process of endochondral ossification in mice by non-enzymatic repression of the activity of the RUNX2 transcription factor (TF) and to control cardiac metabolism in physiological stress situations. In this study, we examined the function of somatic HDAC4-902 frameshift (fs) mutations that are frequently observed in gastric and colon adenocarcinoma patients. Whether these mutations lead to a gain- or a loss-of-function is currently unknown. Here we generated a murine model bearing a germline HDAC4-methionine (M) amino acid (AA) 902-to-histidine (H) frameshift (M902Hfs) mutation. HDAC4-M902Hfs mice phenocopied HDAC4 null mice and present with premature ossification and early postnatal death. Mechanistically, we found that the HDAC4-M902Hfs mutation induced nonsense-mediated mRNA decay, resulting in loss of HDAC4 protein. This loss-of-function (LOF) effect was further supported by increased mRNA and protein expression of runt-related transcription factor-2 (RUNX2) and reduced class IIa HDAC enzymatic activity, indicating that HDAC4 contributes significantly to endogenous class IIa HDAC activity. Patient-derived data suggest that the HDAC4-902fs mutation is associated with reduced mRNA expression of HDAC4. In conclusion, our study identify that HDAC4-902fs mutation is a loss-of-function mutation, but raises the new question whether the loss of non-enzymatic mechanisms or the reduction in class IIa HDAC activity contributes to tumor progression.

## Introduction

During embryonic development, most long bones are formed via endochondral ossification, a process whereby mesenchymal cells differentiate into chondrocytes to form a cartilaginous template during ossification (1). Ossification occurs when chondrocytes in the cartilaginous template undergo hypertrophy, resulting in calcification of cartilage, forming an extensive vascular network and extracellular matrix to produce primary centers of ossification (1). Histone deacetylases (HDACs) regulate developmental gene programs in the vertebrate skeleton by catalyzing the removal of acetyl groups from both nuclear histones and non-histone proteins, including transcription factors (TFs), ultimately leading to closed chromatin conformation and consequently suppression of gene activity (2, 3). HDAC4 is a member of class IIa HDACs (HDAC4, HDAC5, HDAC7 and HDAC9) that is robustly expressed in osteoblasts and prehypertrophic chondrocytes (4). Previous research has revealed that HDAC4 inhibits the action of TFs such as RUNX2 and myocyte-specific enhancer factor 2 (MEF2A, MEF2C and MEF2D) most likely through a non-enzymatic mechanism to suppress chondrocyte hypertrophy (4–9). HDAC4 gene disruption via global knockout in mice promotes premature ossification, restricted neonatal growth and premature demise (< 2-3 weeks) (4). The phenotype observed in HDAC4^-/-^ mice is caused by early chondrocyte hypertrophy, which limits rib cage extension and results in early respiratory failure. The development of cartilaginous templates of endochondral bones occurs normally in HDAC4^-/-^ mice, but the initiation of chondrocyte hypertrophy is hastened, resulting in ectopic bone formation and an increase in RUNX2 activity (1).

Growing evidence has supported that epigenetic enzymes are mutated in various malignancies (10, 11), many of these mutations include nonsense mutations, frameshift mutations, and mutations that cause alternative splicing events, result in premature termination codons (PTCs), and produce frameshifts that generate a downstream stop codon. These mutations are often considered to be loss-of-function (LOF) as PTC-containing transcripts can be degraded by an mRNA surveillance mechanism that termed as nonsense-mediated mRNA decay (NMD) (12–15). NMD activation results in degradation of the transcripts bearing PTCs and no protein will be produced(16–20). In the present study, we used *in vivo* model to investigate the function of a somatic mutation of HDAC4-902fs found in gastric and colon adenocarcinoma patients.

## Results

### The deacetylase domain of HDAC4 is mutated in cancer patients

The HDAC4-902 frameshift (fs) mutation has been described in cancer patients as a somatic mutation (28–31). Two distinct variants of this mutation have been identified: 1) an insertion of the ‘cytosine’ (C) nucleotide at 902 amino acid position leads to a truncated protein at 906 amino acid and has been found in 7 cancer patients changing the codon from Methionine (M) to Histidine (H) (Fig. 1A and Fig. S1); 2) a deletion of the ‘adenine’(A) nucleotide at 902 amino acid position leads to a truncated protein at 956 amino acid and has been found in 20 cancer patients, changing the codon from Methionine (M) to Tryptophan (W) (32–35) (Fig. 1A and Fig. S1). Both variants occur in heterozygous state (28). To characterize their functional impact, we generated a, stable cell lines expressing full-length HDAC4 and two mutants of HDAC4-902fs with an N-terminal Flag tag (Fig. 1B-C). Prior research demonstrated that HDAC3 coimmunoprecipitated with HDAC4 (36) to repress the target genes of HDAC4 (37). Thus, we examined the ability of wild-type HDAC4 (HDAC4-WT) and HDAC4-902fs mutants to interact with HDAC3. To confirm this, we overexpressed Flag-HDAC4-WT versus mutants of HDAC4-902fs in HEK293 cells and performed HDAC activity assay and co-immunoprecipitation (co-IP). Our co-IP data revealed that HDAC4-WT binds strongly to HDAC3, whereas the mutants lost the interaction with HDAC3 and exhibited low HDAC activity towards class I and IIa HDAC substrates (38) (Fig. 1D-E). To determine whether frameshift mutations affected transcriptional repression, we performed luciferase reporter assay using MEF2-responsive elements. As expected, overexpression of Flag-HDAC4 and HDAC4 3S/A (S246A;S467A;S632A) mutants significantly repressed MEF2 activation; by contrast, frameshift mutations failed to repress MEF2C transcription factor (TFs) (Fig. 1F). Taken together, these findings support that HDAC4-902fs mutations disrupt protein deacetylase activity and interaction with HDAC3, and attenuate inhibition of MEF2, which has been shown to occur independent of the enzymatic domain (39, 40).

**Figure 1:**
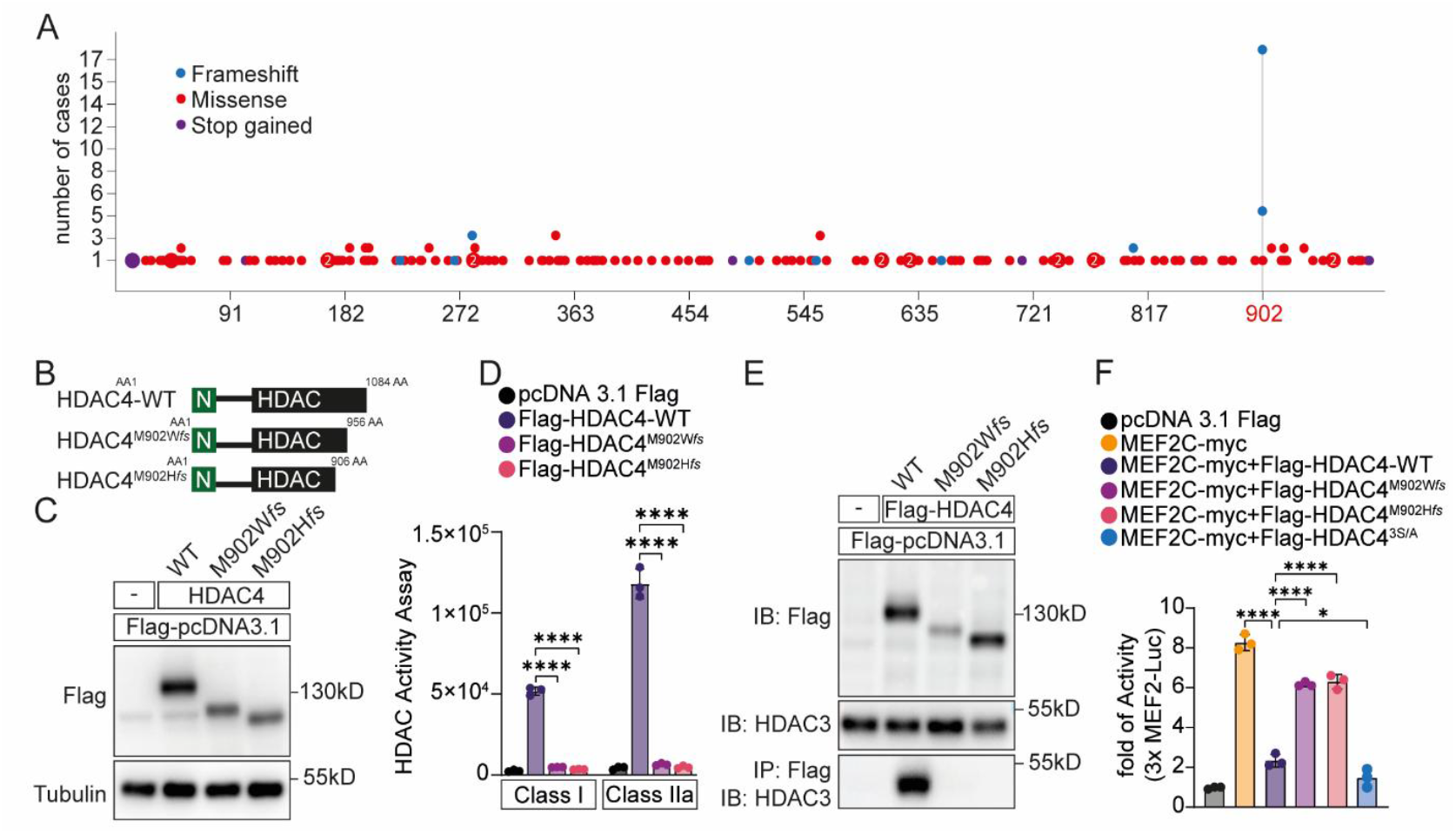
Identification of HDAC4-902 frameshift mutation in Cancer Patients. (A) Somatic mutations of HDAC4 identified in cancer patients were retrieved from cBioPortal. (B) Schematic representation of HDAC4 protein. (C) HEK293T cells were transduced with plenti puro Flag HDAC4-WT and mutants constructs. Empty vector plenti puro Flag was used a negative control. Whole cell-lysates were subjected to immunoblotting with the indicated antibodies. Tubulin was used as a loading control. (D) Class I and Class IIa HDAC activities were measured, 24 hours post-transfection with pcDNA3.1 Flag HDAC4-WT and mutants in HEK293A. (E) HEK293A cells were transfected with pcDNA3.1 Flag, Flag HDAC4-WT and mutants constructs. Lysates were immunoprecipitated with Anti-Flag M2 agarose and immunoblotted with the indicated antibodies. (F) Luciferase assay was performed in HEK293A cells by co-transfecting following plasmids; MEF2c-myc, HDAC4-Flag and mutants constructs. 3xMEF construct, presenting three binding sites for MEF2, was used as positive control. Data were normalized by co-transfecting pRenilla and expressed as mean ± S.D., *n* = 3. Throughout, data are mean ± s.d. *P < 0.05 and **P < 0.01; n.s., P > 0.05; by ANOVA.

### HDAC4^M894H^ mice phenocopy HDAC4 null mice

To investigate the function of the HDAC4-902fs mutation *in vivo*, we generated a knock-in (KI) mouse model of the HDAC4-M902Hfs mutation using CRISPR/Cas9 technology (Fig. 2A). This mutation in mice causes insertion of the ‘Guanine’(G) nucleotide at 894 amino acid (the human counterpart of amino acid 902), changing the codon from Methionine (M) to Histidine (H) and results in a premature stop codon at amino acid 898. Genotyping confirmed the presence of wildtype (WT), heterozygous (Het) and KI alleles (Fig. 2B), which was further mice validated by sanger sequencing (Fig. 2C). HDAC4^M894H^ Het mice were backcrossed with C57BL6/N-WT mice for six generations before used for the current study. Phenotypic analysis revealed that HDAC4^M894H^ (KI) mice displayed a “dome-shaped” heads and misshaped spines similar to HDAC4 null mice within the first three days after birth, which is likely due to disinhibition of RUNX2 TFs in chondrocytes (Fig. 2D-E) (4). Consistent with this severe phenotype, none of the KI mice survived until the weaning period and were smaller relative to WT mice at postnatal days 3, 5, 8 and 12 (Fig. 2F-H). By contrast, HDAC4^M894H^ Het mice survived until adulthood and were morphologically indistinguishable from wild-type littermates.

**Figure 2:**
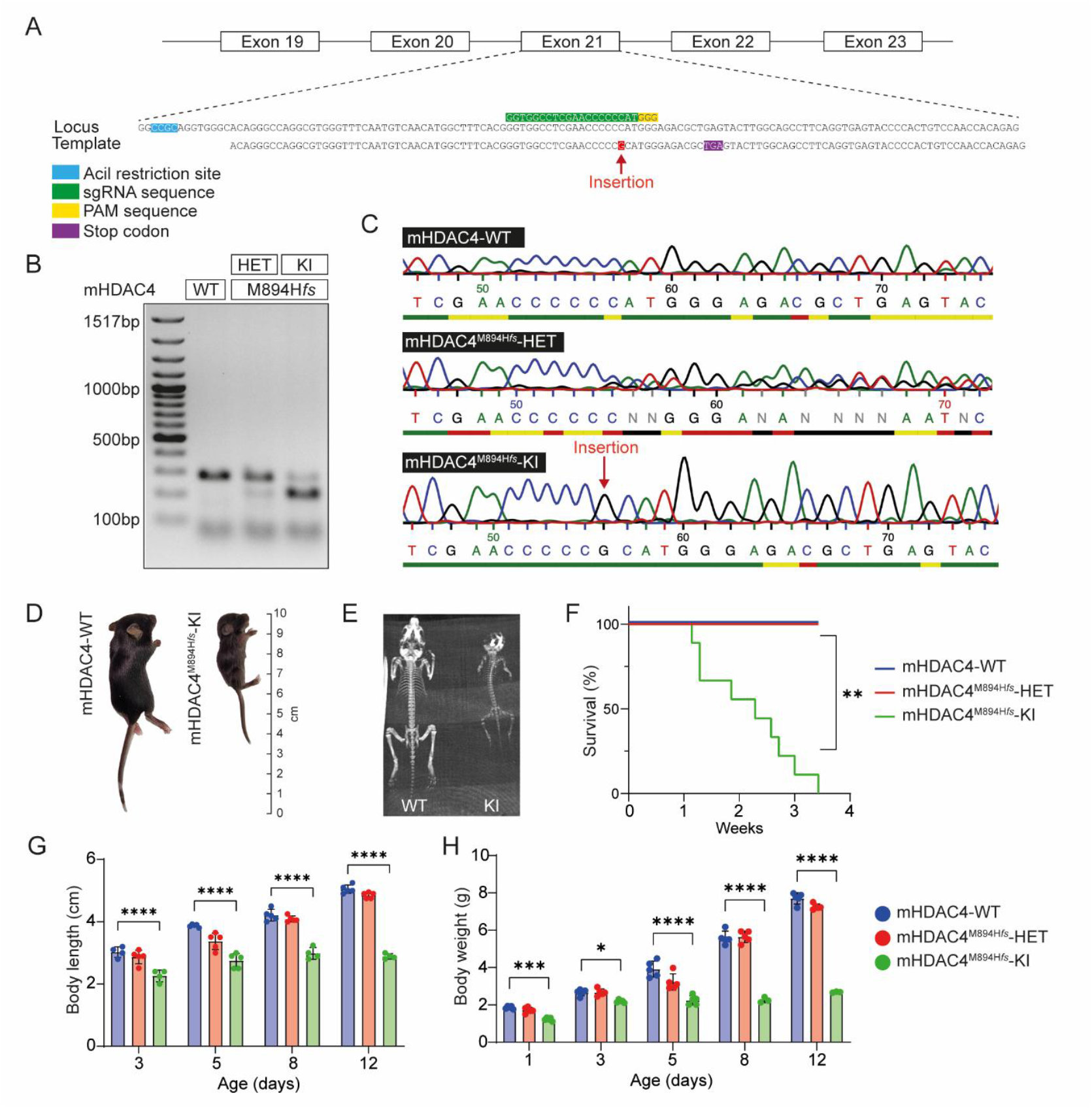
Characterization of CRISPR-generated-HDAC4^M894H*fs*-KI^ mice. (A) A strategy to generate a HDAC4^M894H*fs*^ (KI) mouse line. A sgRNA (green) targeting the respective locus were designed using CRISPOR. A 131 base-pair (bp) template single stranded oligodeoxynucleotide (ssODN) was designed providing not only the insertion guanine (G) nucleotide but also an AciI (blue) restriction site for genotyping purposes. (B) PCR products amplified from genomic DNA were digested with the respective enzymes and fragments were analysed by 2% agarose gel electrophoresis. (C) Representative Sanger sequencing chromatogram showing insertion of G nucleotide in KI mice (red arrow). (D-E) Photographic and CT Scan images of HDAC4-WT mice versus HDAC4^M894H*fs*-KI^ mice at postnatal day 12 age. (F) Kaplan-Meier survival study shows that HDAC4^M894H*fs*-KI^ mice die prior weaning period, whereas heterozygous (HET) mice is indistinguishable from wild type (WT) mice (n=9 per group). (G-H) Body length and body weight of WT, heterozygous and homozygous mice at indicated time intervals (n=4-5 per group). Throughout, data are mean ± s.d. *P < 0.05 and **P < 0.01; n.s., P > 0.05; by ANOVA.

### HDAC4^M894H^ frameshift mutation affects protein and mRNA levels

Western blot and RT-PCR quantification unmasked the absence of HDAC4 protein and transcript in ribs of KI mice (Fig. 3A and Fig. S2A). Notably, the recently reported proteolytic fragment of HDAC4, named HDAC4-NT (51) is also detectable in ribs, raising the question whether this fragment also exerts a function in bone. HDAC4 was also absent in others tissue types including the brain, heart, and skin, where HDAC4 is typically expressed (Fig. S2B). Consistent with reports from the Partidge lab, which have shown that HDAC4 null mice display elevated expression of matrix metalloproteinase-13 (MMP-13) mRNA in the hypertrophic chondrocytes, we found increased expression of MMP-13 in KI mice (Fig. 3B) (41). Furthermore, both protein and mRNA levels of RUNX2 and HDAC5 were elevated in KI mice, indicating a compensatory feedback loop to HDAC5 as described previously (Fig.3A) (42). To examine how HDAC5 upregulation affects rib HDAC activity, we measured total HDAC activity in the ribs. Loss of HDAC4 led to a significant reduction in class IIa HDAC activity in ribs (Fig.3D), indicating that HDAC4 normally contributes a major portion of this enzymatic function. Although HDAC5 expression is upregulated at both the mRNA and protein levels as a compensatory response, this increase is insufficient to fully restore HDAC4 function. This suggests that while HDAC5 may partially compensate for the absence of HDAC4, the overall deacetylase activity of the class IIa family remains lower than in wild-type tissue. Thus, the elevated HDAC5 represents a feedback mechanism attempting to balance the loss of HDAC4, but it cannot completely substitute for HDAC4’s functional contribution in the ribs. Moreover, we found an increase of total H3-acetylation in the mutant mice, suggesting that HDAC4 may in fact act as a histone deacetylase or dislocates HDAC3 from chromatin regions (Fig.3A). The specific histone residues involved remain to be identified. These results indicate that the somatic mutation of HDAC4^M894Hfs^ *in vivo* phenocopies HDAC4 null mice by LOF.

**Figure 3:**
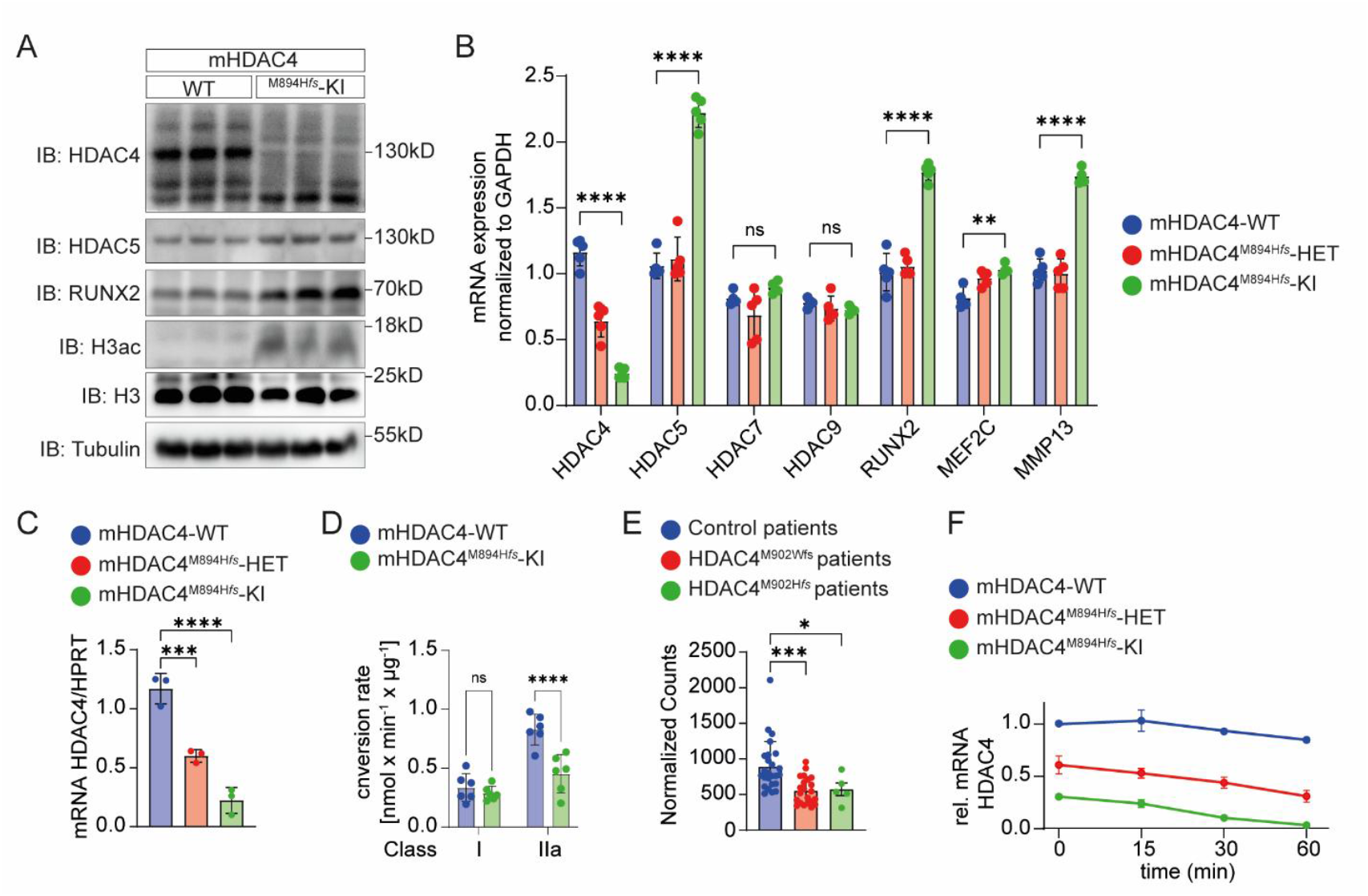
HDAC4^M894H*fs*^ mutation affects HDAC4 protein and mRNA stability. (A) Ribs from WT and KI mice were lysed and subjected to immunoblotting with the indicated antibodies. Mutant mice showed absence of HDAC4 protein,increased HDAC5 and acetylation of Histone H3. Tubulin and histone H3 was used as a loading control (B) qRT – PCR analysis of rib cartilage showed upregulation of HDAC5, RUNX2, MEF2C and MMP-13 in mutant mice, classical genes associated with endochondrial ossification. (C) qRT –PCR of MEFs (E12.5) derived from KI mice showed less HDAC4 mRNA expression to WT controls. (D) RNA-seq data from adenocarcinoma patients demonstrated that cases carrying an HDAC4-902 frameshift mutation showed reduced HDAC4 transcript levels compared to patients without the mutation. (E) MEF cells derived from WT, HET and KI mice were treated with the RNA polymerase inhibitor ActinomycinD (ActD) and RNA was extracted at various time intervals and HDAC4 mRNA levels were analyzed by qRT–PCR. 18S was used as the housekeeping gene. Data are presented as mean ± s.d. *P < 0.05; \**P < 0*.*01;* n.s., P > 0.05; one-way ANOVA.

### Analysis of RNA-Seq data from HDAC4-902fs adenocarcinoma patients showed less HDAC4 transcript

Given that KI model showed complete loss of HDAC4 protein and transcript, therefore we hypothesized that patients with these mutations would have less HDAC4 transcript. To answer this question, we analyzed RNA-seq data from 21 patients with HDAC4-M902Wfs and 5 patients from HDAC4-M902Hfs from The Cancer Genome Atlas (TCGA) database. As a control, we included cancer patients with the same sex, age, and race who did not carry HDAC4 mutations. Remarkably, patients with HDAC4-902fs mutations had a significantly lower HDAC4 transcript level compared to cancer patients without HDAC4 mutations (Fig. 3E).

### Nonsense-mediated mRNA decay (NMD) activation in HDAC4^M894H^ mice degrades HDAC4 Mrna

Our *in vivo* data from KI mice and RNA-Seq data from HDAC4-902fs cancer patients collectively support that this frameshift mutation produced unstable HDAC4 mRNA, and this mRNA could be degraded *in vivo* by NMD, which recognizes and degrades mRNA harboring premature termination codon (PTC). Therefore, to test this hypothesis, we measured mRNA decay in murine embryonic fibroblasts (MEF) cells derived from pregnant female KI mice (Fig. 3C). MEF cells were treated with the RNA polymerase inhibitor ActinomycinD (ActD) and RNA was extracted at various time intervals, and then HDAC4 mRNA levels were analyzed by real-time PCR. For comparison we also treated WT and Het MEF cells with ActD. HDAC4 mRNA levels were stable in WT-MEF cells for at least 60 mins following ActD treatment, whereas in Het and KI-MEF cells HDAC4 mRNA levels were significantly lower after 15mins treatment (Fig. 3F). Taken together, these data suggest that HDAC4 mRNA is rapidly degraded in KI mouse and blocking mRNA degradation could improve the stability of mutant HDAC4 mRNA.

### NMDi inhibited NMD activation in HDAC4^M894H*fs*-KI^ KI MEF cells and enhanced protein stability

NMD inhibition can result in the upregulation of non-mutated, mutated and alternatively spliced mRNAs (43). To test this, we used the two putative NMD inhibitors (NMDIs) Amlexanox (44) and NMDI14 (45) and measured NMD inhibition by assessing HDAC4 mRNA levels from WT, Het and KI isolated MEF cells. Treatment for 6 hours with the varying doses of NMDI14, resulted in an increase of HDAC4 mRNA in Het and KI MEF cells (Fig.S3). Particularly Amlexanox was more effective in preventing the mRNA degradation of HDAC4, which resulted in a fourfold increase in HDAC4 mRNA expression in the KI MEF cells (Fig. 4A). When Flag-HDAC4-902fs mutants were transduced *in vitro*, we observed less protein expression as compared to HDAC4-WT (Fig.1C). To investigate if Amlexanox can enhance protein stability, we treated HEK293A cells with Amlexanox for 6 hours after transfecting Flag-HDAC4 WT and Flag-HDAC4-M902Hfs mutation. After extracting RNA and protein from the cells, we observed that Amlexanox increased protein and mRNA stability in the HDAC4-M902Hfs mutant (Fig.4B-C). Taken together our data suggest that pharmacological inhibition of NMD increased the mRNA and protein stability of HDAC4-M902Hfs by preventing degradation of HDAC4.

**Figure 4:**
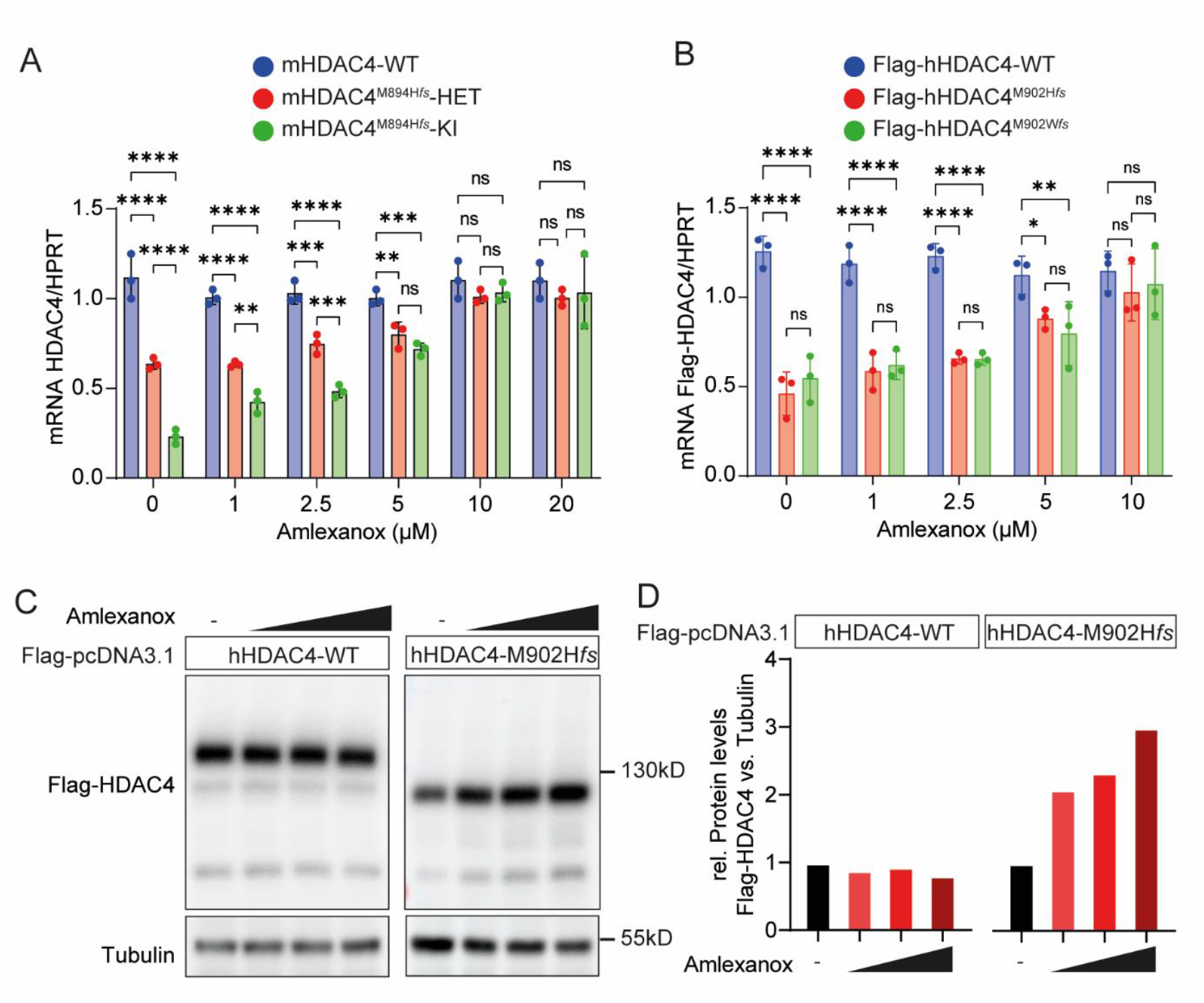
Nonsense-mediated mRNA decay (NMD) inhibition restores mRNA and protein stability in Mutant. (A) qRT–PCR analysis of MEFs derived from WT, Het and KI mice, showing that treatment with NMD inhibitor Amlexanox increased HDAC4 mRNA levels. (B) qRT–PCR analysis of Flag HDAC4-WT and mutants constructs overexpressed in HEK293A cells following treatment with NMD inhibitors. HDAC4 mRNA stability was assessed using primers spanning the (forward) Flag sequence and (reverse) HDAC4 exon 2. (C) Flag HDAC4 and mutant M902H*fs* were transfected in HEK293A cells and treated *with NMD inhibitor A*mlexanox. Immunoblotting analysis revealed increased protein stability of the mutant upon NMD inhibitor. Tubulin was used as a loading control. (D) Quantification of protein band intensities corresponding to panel C. Data are presented as mean ± s.d. *P < 0.05; \**P < 0*.*01; n*.*s*., *P > 0*.*05; one-way ANOVA*

## Discussion

Histone deacetylases (HDACs) control acetylation by removing acetyl groups from lysine residues of histone and nonhistone proteins, resulting in closed chromatin condensation and transcriptional suppression. Mutations in HDAC genes are linked to tumor progression due to aberrant transcription of essential genes that regulate important biological activities such as cell proliferation, cell cycle regulation and apoptosis. Mutations in HDACs have been identified in a number of cancers, with somatic mutation of HDAC1 found within 8% of dedifferentiated human liposarcomas, while the loss-of-function of HDAC2 caused by a frameshift mutation was studied in human epithelial and colorectal malignancies (46). The current study aimed to study the functional impact of an HDAC4-902*fs* mutation on the pathogenesis of somatic cancers and its potential role in the pathogenesis of these malignant tissues. HDAC4-902*fs* is the only mutation detected in multiple cancer patients, and it occurs in the deacetylase domain of HDAC4. Although this mutation has been discovered in a number of cancer patients, its prevalence, origins and role remain mysterious. In vertebrates, class IIa HDACs were reported to be catalytically inactive epigenetic readers quickly recruited on H3K27ac loci (47, 48). Here they scrutinize the acetylation status of H3K27ac through forming a class IIa corepressor complex with nuclear receptor corepressor 1 (NCoR1) or its homolog silencing mediator of retinoic and thyroid receptors (SMRT) and HDAC3 to repress gene expression (37). Any mutations on the deacetylase domain leading to the suppression of HDAC4 binding to NCoR1/SMRT and to HDAC3 result in the loss of HDAC4 deacetylase activity. We identified that HDAC4-902fs mutants doesn’t interact with HDAC3 and showed reduced class I and class IIa HDAC enzymatic activity.

CRISPR/Cas9 generation of HDAC4-M902Hfs (KI) mice phenocopied HDAC4 null mice by displaying “dome-shaped” heads and misshaped spines resulting in disinhibition of RUNX2 TFs. With this mutation, we found that KI mice lack HDAC4 mRNA and protein. None of the KI mice survived until weaning, while the majority of mice died before postnatal day 12. NMD is a translation-dependent mRNA surveillance mechanism in eukaryotes that helps preserve the quality of gene expression. NMD causes rapid decay of aberrant mRNAs with a premature termination codon (PTC) (16, 49) and preventing the generation of truncated proteins. MEF cells from KI mice, when treated with NMDI increased the HDAC4 mRNA levels similar to WT. These findings show presence of PTC on HDAC4-902 promotes NMD, and inhibiting NMD reduces the degradation of HDAC4 mRNA. Thus, these data cannot answer the question whether intrinsic enzymatic activity of HDAC4 contributes to cancer progression. Instead loss of function affects both enzymatic and N-terminal mediated non-enzymatic actions of HDAC4. What are the implications of the data derived from HDAC4^M894H*fs*-KI^ mice for cancer? Using publicly available data we found that patients with GI-related adenocarcinoma harboring with the somatic HDAC4-902fs mutations had lower HDAC4 transcript levels, which is similar to the phenotype of KI mice reported in the current study. The lethality of the germline mutation in mice is unfortunately limiting us from recapitulating the human observations in a mouse model to demonstrate a potential role in tumor progression. It could be hypothesized that e.g. radiotherapy induces HDAC4-902fs mutations that then contribute to protection. Support comes from efforts that led to the development of enzymatic class IIa HDAC inhibitors as immunomodulators to reduce tumor malignancy (50). Although HDAC5 expression is upregulated in response to HDAC4 loss, overall class IIa HDAC activity remains reduced, indicating that HDAC5 cannot fully compensate for HDAC4 enzymatic function. This highlights that therapies targeting class IIa HDACs may still be effective even in the context of HDAC5 compensation.

The potential role of HDAC4-902fs as a protective somatic mutation in cancer warrants further exploration. Reduced HDAC4 activity may limit oncogenic transcriptional programs, and pharmacologic inhibition or genetic loss of HDAC4 has been shown to restrict tumor progression in other models (47). Future studies are needed to further investigate how HDAC4-902fs affects cancer cell proliferation, therapeutic response, and immune regulation. However, whether the reduction of enzymatic activity or the loss of the entire protein including its non-enzymatic mechanisms is functionally relevant remains unanswered by the use of the model reported here. Specific mutations in the active site in *in vivo* models may answer this clinically relevant question and may support the need for the development of class IIa HDAC inhibitors which have not yet reached clinical testing.

In conclusion, our study demonstrates that the HDAC4-902fs mutation results in NMD-mediated transcript degradation, loss of HDAC4 protein, and reduced class IIa HDAC activity. These findings provide mechanistic insight into how HDAC4-902fs mutations contribute to functional LOF and highlight HDAC4 loss as a potential epigenetic vulnerability in human cancer. However, more optimized models are needed to differentiate between the enzymatic and non-enzymatic actions of HDAC4 in tumor models.

## Methods

### Sex as a biological variable

Our study examined male and female animals and similar findings were identified in both sexes.

### Animal Experiments

All animal experimental work was approved by the Institutional Animal Care and Use Committee at the Regierungspräsidium Karlsruhe, Germany. HDAC4-^M894H*fs*^ knockin (KI) mice were generated using CRISPR/Cas9 technology (21). To generate potential sgRNA sequences, the genomic region around the targeting site was entered and the mouse genome GRCm38/mm10 (UCSC Dec. 2011) and the protospacer adjacent motif (PAM) NGG from Streptococcus pyogenes were selected. Following sgRNAs sequence were chosen around the targeting site by using CRISPOR: GGTGGCCTCGAACCCCCCAT*GGG*. T7 strings coding for sgRNAs were commercially provided by Thermo Scientific, which were used for in vitro transcription with Mega Short Script T7 Kit (Thermo Scientific) and Mega Clear Kit (Thermo Scientific) according to the manufacturer’s protocol. Injection Mix containing 3µM of a 131 bp template (5’-acagggccaggcgtgggtttcaatgtcaacatggctttcacgggtggcctcgaaccccccatgggagacgctgagtacttggcagccttcaggtgagtaccccactgtccaaccacagaggctggttcttc-3’, Eurofins Genomics, Germany), 25ng/µL of in vitro transcribed sgRNA, 25ng/µL Cas9 mRNA (Thermo Scientific) and 25ng/µL Cas9 protein (New England-Biolabs) diluted in sterile-filtered injection buffer (5mM Tris, 0.1mM EDTA, pH 7.4) were injected into the cytoplasm of zygotes from super ovulated female C57BL6/N mice at Interfaculty Biomedical Facility (IBF) Heidelberg University Germany. Later zygotes were implanted into foster mice. Positive founder mice were back crossed to C57BL6/N wildtype mice for six generations before they were used for experiments. C57BL/6N mice for breeding were ordered from Janvier Labs (Le Genest-Saint-Isle, France) All the animals were housed in a specific pathogen–free environment with a 12-hour light/12-hour dark cycle and provided with water and standard food ad libitum.

### Genotyping protocol for HDAC4^M894H*fs*^ knockin mice

DNA was extracted from tissue biopsies using the DNeasy Blood & Tissue Kit (Qiagen #69506, Germany) according to the manufacturer’s protocol for DNA isolation from tissue. In order to differentiate between wild type and the mutant allele, an additional restriction enzyme (AciI) (NEB #R0551S) was introduced in the sequence, and this helps to identify the mutant allele upon restriction digest of the PCR product. After the restriction digestion with AciI, DNA samples were purified and sequenced by Sanger method.

### Plasmid construction

The following plasmids were used: pcDNA3.1 MEF2c-myc, pcDNA3.1 Flag HDAC4 3S/A (6). HDAC4 was subcloned into pcDNA3.1 Flag to have a N terminal tag, pcDNA3.1 Flag HDAC4 M902H*fs* and M902Wfs mutants were generated by overlap extension using Phusion® High Fidelity DNA Polymerase (NEB #M0530S). plenti puro HDAC4 M902H*fs (*was gift from Claudio Brancolini’s laboratory).

### Murine Embryonic Fibroblasts (MEF) cell isolation

MEF cell isolation was performed as previously described (22).

### Cell culture, transfections, and reagents

Dulbecco’s Modified Eagle Medium (DMEM) with 10% Fetal Bovine Serum (FBS) and 1% Penicillin-Streptomycin was used to cultivate Murine Embryonic Fibroblasts (MEF) and Human Embryonic Kidney 293A/T cells (HEK293A/T), respectively. plenti puro HDAC4 and mutants of HDAC4-902fs were transfected with Lipo2000. pcDNA3.1 Flag HDAC4 and mutants of HDAC4-902*fs* were transfected according to Gene jammer transfection protocol (Agilent #204130). Retroviral infections were as previously described (23). To inhibit Nonsense mediated mRNA decay (NMD), inhibitors were purchased from MedChemExpress (Amlexanox #HY-B0713, NMDi14 #HY-111374).

### RNA Isolation and Reverse transcription-quantitative PCR (qPCR)

Total RNA was extracted with TRIzol reagent and 500∼1000ng of RNA was reverse transcribed to cDNA using the QuantiTect RT kit (Qiagen #205311). Gene expression levels were performed by qPCR analyses by using LightCycler 480 SYBR Green I Master (Roche #04887352001). Relative quantification was performed to evaluate target genes expression by normalizing to GAPDH.

### Western Blotting, Immunoprecipitation and antibodies

Total tissue protein lysates from different organs (ribs, skin, heart and brain) were lysed in Radio-Immunoprecipitation Assay (RIPA) buffer and proteins were separated on Tris-Glycine SDS-PAGE and transferred electrophoretically onto nitrocellulose membrane in Pierce® Western Blot Transfer Buffer (Thermo Scientific, Waltham, Massachusetts). After blotting, membranes were rinsed with TBS-T (100 mM Tris, pH 8.0, 1500 mM NaCl, 0.5% (v/v) Tween-20) for 30 minutes and incubated in primary antibody overnight at 4°C with shaking. Primary antibodies are as follows: Anti-Flag Antibody (Thermo Fisher Scientific #MA1-91878), GAPDH (Merck Millipore #MAB374), HDAC4 (abcam #12172), HDAC4 (abcam #123513), H3-ac (active motif #39140), H3 total (abcam #1791), HDAC5 (Cell Signaling Technology #98329), RUNX2 (Cell Signaling Technology #12556) and HDAC3 (Cell Signaling Technology #85057). For Immunoprecipitation (IP), protocol from (PMID-33966634), was followed.**mRNA stability assay**. MEF cells were treated with 2.5 μM Actinomycin D (ActD) (Sigma-Aldrich) and cells were harvested at different time points 15, 30, and 60 min. Total RNA was extracted and analysed by real-time PCR; 18S was used as the housekeeping gene in these experiments (24, 25).

### HDAC activity assay

HDAC activity assay were performed as previously described by Lemon et al. (26).

### RNA-sequencing analysis

Transcriptomic data from human malignancies were obtained from The Cancer Genome Atlas (TCGA) based on DNA sequencing analysis of the HDAC4 gene. Details regarding computational tools and coding scripts required in the current study are provided as an online supplement via GitHub data repository: https://github.com/mepepin/HDAC4_Onc. Alignment was accomplished using *STAR* (2.7.10a), with differential gene expression calculated using *DESeq2* (1.32.0) within the R (4.1.2) statistical computing environment. In this way, we estimated dispersion via maximum-likelihood and minimized it using the empirical Bayesian method to provide normalized counts based on dispersion estimates and sample size. We then calculated differential expression from normalized read counts via Wald test with Bonferroni *post hoc* adjusted *P-*value for each aligned and annotated gene. Statistical significance was assumed as unpaired two-tailed Bonferroni-adjusted *P* < 0.05, with biological significance assumed when |Fold-Change| > 50% with normalized count sum > 1.1 (27).

### Statistics

For all pairwise comparisons, the Shapiro-Wilk test for normality was performed to determine the most appropriate statistical test. All pairwise factors exhibiting a parametric distribution were evaluated using student t-test with Benjamini-Hochberg adjustment; otherwise, a Mann-Whitney test was used. All data are reported as mean ± standard deviation unless otherwise specified. Statistical analyses and data visualization were completed using GraphPad Prism version 9.3.1 for Macintosh (GraphPad Software, San Diego, CA) and R software, version 4.1.2 (R Foundation for Statistical Computing, Vienna, Austria). Statistical significance was assigned when *P* < 0.05.

## Acknowledgments

We would like to thank you Prof.Claudio Brancolini and Prof. Eros Di Giorgio for sharing the plasmid PWZL-Hygro HDAC4-M902Hfs. We thank Walter Mier for assistance with CT scan and Dr. Shanmukha Doddi for technical assistance.

