## Supplementary Figures for "Somatic Hdac4-902fs mutations lead to loss of HDAC4 function through nonsense-mediated mRNA degradation"

### Supplemental Figures

**HDAC4-WT**

| | | | | | | | | | | | | | | | | | | | |
|---|---|---|---|---|---|---|---|---|---|---|---|---|---|---|---|---|---|---|---|
| 901 | 902 | 903 | 904 | 905 | 906 | 907 | 908 | 909 | 910 | 911 | 912 | 913 | 914 | 915 | 916 | 917 | 918 | 919 | 920 |
| CCC | **ATG** | GGA | GAC | GCT | GAG | TAC | TTG | GCG | GCC | TTC | AGA | ACG | GTG | GTC | ATG | CCG | ATC | GCC | AGC |
| P | M | G | D | A | E | Y | L | A | A | F | R | T | V | V | M | P | I | A | S |
| 921 | 922 | 923 | 924 | 925 | 926 | 927 | 928 | 929 | 930 | 931 | 932 | 933 | 934 | 935 | 936 | 937 | 938 | 999 | 940 |
| GAG | TTT | GCC | CCG | GAT | GTG | GTG | CTG | GTG | TCA | TCA | GGC | TTC | GAT | GCC | GTG | GAG | GGC | CAC | CCC |
| E | F | A | P | D | V | V | L | V | S | S | G | F | D | A | V | E | G | H | P |
| 941 | 942 | 943 | 944 | 945 | 946 | 947 | 948 | 949 | 950 | 951 | 952 | 953 | 954 | 955 | 956 | 957 | | | 1084 |
| ACC | CCT | CTT | GGG | GGC | TAC | AAC | CTC | TCC | GCC | AGA | TGC | TTC | GGG | TAC | CTG | ACG | ... | ... | TGA |
| T | P | L | G | G | Y | N | L | S | A | R | C | F | G | Y | L | T | | | * |

**HDAC4-M902H*fs***

| | | | | | |
|---|---|---|---|---|---|
| 901 | 902 | 903 | 904 | 905 | 906 |
| CCC | CAT | GGG | AGA | CGC | TGA |
| P | H | G | R | R | * |

**HDAC4-M902W*fs***

| | | | | | | | | | | | | | | | | | | | |
|---|---|---|---|---|---|---|---|---|---|---|---|---|---|---|---|---|---|---|---|
| 901 | 902 | 903 | 904 | 905 | 906 | 907 | 908 | 909 | 910 | 911 | 912 | 913 | 914 | 915 | 916 | 917 | 918 | 919 | 920 |
| CCC | TGG | GAG | ACG | CTG | AGT | ACT | TGG | CGG | CCT | TCA | GAA | CGG | TGG | TCA | TGC | CGA | TCG | CCA | GCG |
| P | W | E | T | L | S | L | W | R | P | S | E | R | W | S | C | R | S | P | A |
| 921 | 922 | 923 | 924 | 925 | 926 | 927 | 928 | 929 | 930 | 931 | 932 | 933 | 934 | 935 | 936 | 937 | 938 | 999 | 940 |
| AGT | TTG | CCC | CGG | ATG | TGG | TGC | TGG | TGT | CAT | CAG | GCT | TCG | ATG | CCG | TGG | AGG | GCC | ACC | CCA |
| S | L | P | R | M | W | C | W | C | H | Q | A | S | M | P | W | R | A | T | P |
| 941 | 942 | 943 | 944 | 945 | 946 | 947 | 948 | 949 | 950 | 951 | 952 | 953 | 954 | 955 | 956 | 957 | 958 | 959 | 960 |
| CCC | CTC | TTG | GGG | GCT | ACA | ACC | TCT | CCG | CCA | GAT | GCT | TCG | GGT | ACC | TGA | CGA | AGC | AGC | TGA |
| P | L | L | G | A | T | T | S | P | P | A | A | S | G | T | * | R | S | S | * |

**Supplementary Figure S1**

Description and identification of HDAC4-902 somatic mutations in cancer patients. *refers to stop codon.

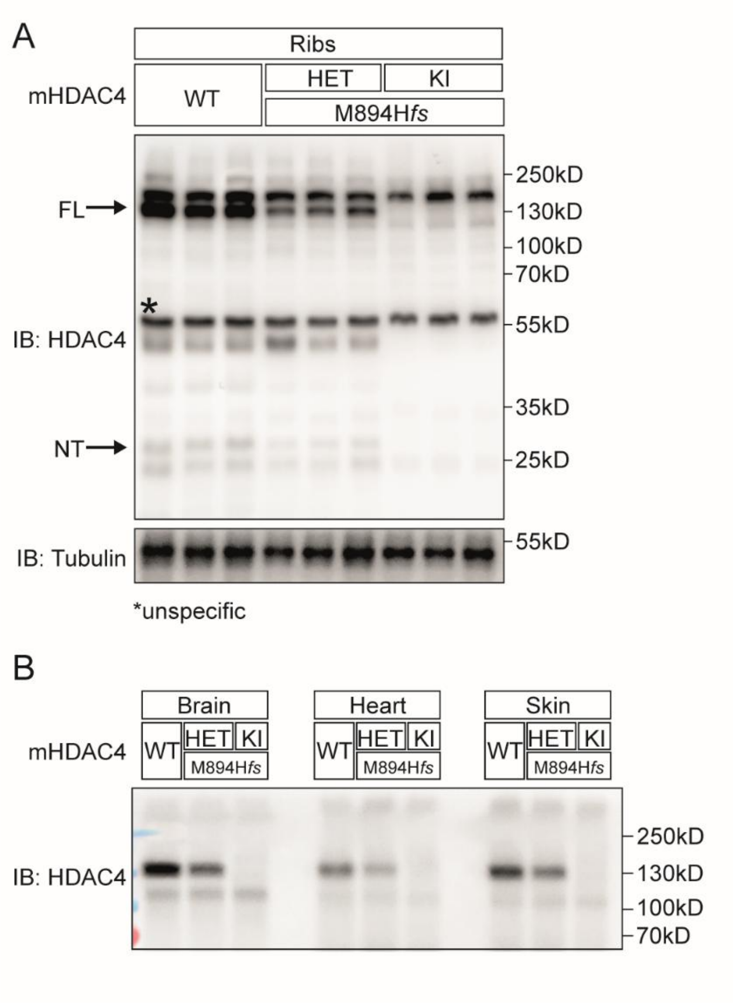


**Supplementary Figure S2**

(A) HDAC4 protein expression in ribs of WT, HET and KI mice. The proteolytic product HDAC4-NT (51) can be detected in ribs. *These bands observed in KI mice seem to be caused by cross reactivity and are unspecific.

(B) HDAC4 protein expression of WT, HET and KI mice in various tissues as indicated.

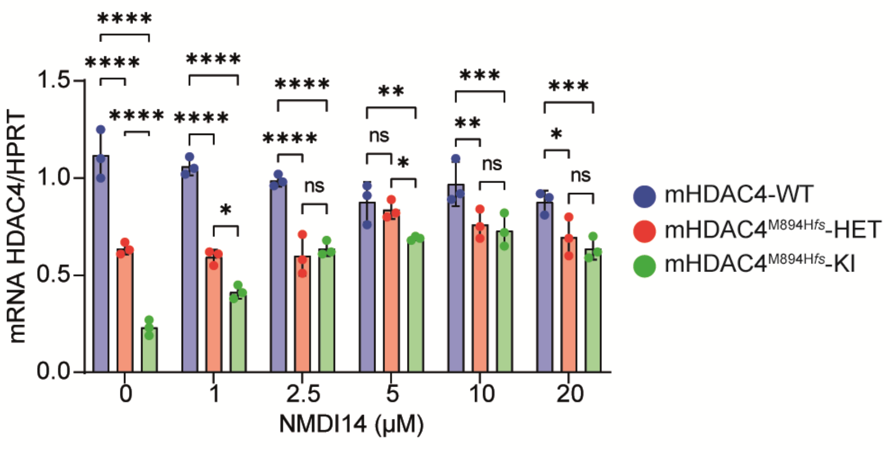


**Supplementary Figure S3**

qRT–PCR analysis of MEFs derived from WT, Het and KI mice, showing that treatment with NMDi-14 inhibitors increased HDAC4 mRNA levels.
